# Bacterial Hsp90 mediates the degradation of aggregation-prone Hsp70-Hsp40 substrates preferentially by HslUV proteolysis

**DOI:** 10.1101/451989

**Authors:** Bruno Fauvet, Andrija Finka, Marie-Pierre Castanié-Cornet, Anne-Marie Cirinesi, Pierre Genevaux, Manfredo Quadroni, Pierre Goloubinoff

**Affiliations:** Department of Plant Molecular Biology, University of Lausanne, CH-1015 Lausanne, Switzerland.; Department of Ecology, Agronomy and Aquaculture, University of Zadar, 23000 Zadar, Croatia.; Laboratoire de Microbiologie et de Génétique Moléculaires, Centre de Biologie Intégrative, Université de Toulouse, CNRS, UPS, 31062 Toulouse Cedex 09, France.; Protein Analysis Facility, University of Lausanne, CH-1015 Lausanne, Switzerland.

**Keywords:** DnaK, DnaJ, HtpG, HslV, proteostasis

## Abstract

Whereas in eukaryotic cells, the Hsp90s are profusely-studied molecular chaperones controlling protein homeostasis together with Hsp70s, in bacteria, the function of Hsp90 (HtpG) and its collaboration with Hsp70 (DnaK) remains unknown. To uncover physiological processes depending on HtpG and DnaK, we performed comparative quantitative proteomic analyses of insoluble and total protein fractions from unstressed wild type *E. coli*, and from knockout mutants *ΔdnaKdnaJ* (ΔKJ), *ΔhtpG* (ΔG) and *ΔdnaKdnaJΔhtpG* (ΔKJG) and compared their growth rates under heat-stress also with *ΔdnaKdnaJΔhslV*. Whereas, expectedly, mutant ΔG showed no proteomic differences with wild-type, ΔKJ expressed more chaperones, proteases and ribosomes and dramatically less metabolic and respiratory enzymes. Unexpectedly, we found that ΔKJG showed higher levels of metabolic and respiratory enzymes and both ΔKJG and *ΔdnaKdnaJΔhslV* grew better at 37^o^C than ΔKJ. The results indicate that bacterial Hsp90 mediates the degradation of aggregation-prone Hsp70-Hsp40 substrates, preferably by the HslUV protease.

**Significance statement:** The molecular chaperones Hsp70 and Hsp90 are among the most abundant and well-conserved proteins in all realms of life, forming together the core of the cellular proteostasis network. In eukaryotes, Hsp90 functions in collaboration with Hsp70; we studied this collaboration in *E. coli*, combining genetic studies with label-free quantitative proteomics in which both protein abundance and protein solubility were quantified. Bacteria lacking Hsp70 (DnaK) and its co-chaperone DnaJ (ΔdnaKdnaJ) grew slower and contained significantly less key metabolic and respiratory enzymes. Unexpectedly, an additional deletion of the Hsp90 *(htpG)* gene partially restored the WT phenotype. Deletion of the HslV protease in the ΔdnaKdnaJ background also improved growth, suggesting that bacterial Hsp90 mediates the degradation of Hsp70 substrates, preferentially through HslV.

At 37^o^C *ΔdnaKdnaJ E. coli* mutants grow slower than wild type cells. Quantitative proteomics shows that compared to wild type cells, *ΔdnaKdnaJ* cells grown at 30^o^C contain significantly less key metabolic and respiratory enzymes. Unexpectedly, deletion of the *HtpG* gene in the Δ *dnaKdnaJ* background ameliorates growth at 37^o^C and partially restores the cellular levels of some metabolic and respiratory enzymes.

## Introduction

Various stresses and mutations increase the propensity of labile cellular proteins to transiently unfold and convert into stably misfolded and aggregated conformers devoid of their dedicated biological functions. Moreover, early forms of protein aggregates may be toxic to eukaryotic cells by causing membrane damage, the release of reactive oxygen species, leading to cell death, aging and degenerative diseases (Lashuel and Lansbury 2006, Hinault, Farina-Henriquez-Cuendet et al. 2011). Various chaperone orthologues from the conserved Hsp70s and Hsp90s families are present at concentrations on the order of tens of micromolar in all ATP-containing compartments of the eukaryotic cells (Finka, Sood et al. 2015). Mass-wise, the main Hsp70s and Hsp90s in human cells are, respectively, HspA8 & HspA1A and Hsp90aa1 & Hsp90ab1 in the cytosol, HspA5 (Grp78, BIP) and Hsp90b1 (Grp94, endoplasmin) in the endoplasmic reticulum and HspA9 (the DnaK homologue) and Trap-1 (TNF receptor associated protein 1, Hsp75, HSP90L, the HtpG homologue) in the mitochondria. Typically assisted in the cytosol by the Hsp70-Hsp90 organizing protein (HOP), both stand at the centre of a network of molecular chaperones, co-chaperones and proteases controlling all aspects of cellular protein homeostasis. They assist *de novo* folding of nascent polypeptides, drive protein translocation, control polypeptide maturation and assembly into functional complexes and finally, they target native and stress-damaged proteins to degradation by the proteasome or by other ATP-dependent chaperone-gated proteases, such as FtsH, Lon, HslUV and ClpXP (Finka, Mattoo et al. 2016, Hartl 2017). Attesting for their centrality under physiological and stress conditions, members of the Hsp90 and Hsp70 families are among the most abundant proteins of the human proteome, accounting for 2-5 % of the total protein mass of some cells (Finka, Sood et al. 2015).

As suggested by their already elevated basal levels in unstressed cells, Hsp90s and Hsp70s also control a plethora of physiological processes, such as the conversion of active protein oligomers into transiently inactive sub-complexes, as in the case of clathrin cages, IκB and HSF1 oligomers (Pratt, Morishima et al. 2010, Finka, Mattoo et al. 2016). In metazoans, Hsp70s and Hsp90s are also involved in the activation of steroid hormone receptors, the transduction of apoptotic signals, the translocation of cytosolic polypeptides into organelles and the targeting by ubiquitination of stress-damaged proteins to the proteasome for controlled degradation (Lackie, Maciejewski et al. 2017). The down-regulation or knockout mutations of Hsp90 genes dramatically affects cellular growth and reduces survival following various stresses, such as heat shock (Franzosa, Albanese et al. 2011). Low amounts of specific Hsp90 inhibitors may induce a mild heat-shock-like response in various eukaryotes, including animals and plants (Finka, Cuendet et al. 2012, Fierro-Monti, Echeverria et al. 2013, Fierro-Monti, Racle et al. 2013, Lee, Gao et al. 2013). Ultimately, Hsp90 inhibitors can induce cell death, rendering Hsp90s attractive targets for cancer therapy (Ayrault, Godeny et al. 2009).

*E. coli* cells express a major form of Hsp70, DnaK, and a single Hsp90, HtpG. Both sequence-and structure-wise, HtpG and DnaK share a very a high degree of homology with their respective eukaryotic Hsp90s and Hsp70s counterparts, strongly suggesting that they carry similar collaborative physiological-and stress-related proteostasis functions in bacteria, mitochondria and chloroplasts. The deletion of the single *dnaK gene* from *E. coli* causes a severe growth phenotype at 37°C and at 42°C, the strong accumulation of insoluble protein aggregates (Mogk, Tomoyasu et al. 1999). SILAC-based comparative proteomic studies showed that at 30°C, *ΔdnaKdnaJ* cells degrade a great number of aggregation-prone Hsp70 substrates than WT cells (Calloni, Chen et al. 2012). In contrast, the deletion of the sole *htpG* gene has virtually no phenotype even at 37°C except for a deficiency in the CRISPR-Cas adaptive immunity (Yosef, Goren et al. 2011); and only at elevated temperatures (42°C to 46°C) where *htpG* mutants exhibit mild growth retardation (Bardwell and Craig 1988, Thomas and Baneyx 1998) and a minor accumulation of aggregated proteins (Thomas and Baneyx 2000). In cyanobacteria, the deletion of *htpG* only results in a mild phycobilisome assembly phenotype under physiological conditions (Sato, Minagawa et al. 2010), but has otherwise little to no measurable impact on their physiology. Yet, *htpG* remains highly conserved and is clearly expressed in most eubacteria thus far investigated, suggesting that a specific essential biological function of HtpG, especially under stress, remains to be identified. Experimental evidence suggests that HtpG functions in close collaboration with DnaK. Using *in vitro* assays with purified chaperones, Genest *et al*. showed that when assisted by GrpE and especially by CbpA, which is a DNAJB type of bacterial Hsp40, HtpG and DnaK collaborate at converting artificially unfolded or misfolded polypeptides into native proteins, in an ATP hydrolysis-dependent manner (Genest, Hoskins et al. 2011). Moreover, this cooperative activity involves the direct interaction between DnaK and HtpG (Nakamoto, Fujita et al. 2014, Genest, Hoskins et al. 2015), with the DnaK-HtpG interface being localized to the DnaJ-binding site within DnaK (Kravats, Doyle et al. 2017). Moreover, the collaboration between Hsp70 and Hsp90 has also been demonstrated in other organisms, such as yeast (Genest, Reidy et al. 2013) and metazoans where Hsp70/Hsp90 interactions are key to the maturation of the steroid hormone receptor (Li, Soroka et al. 2012).

Here, we addressed by label-free quantitative proteomic analysis of total and insoluble protein fractions from WT, *ΔdnaKJ* (ΔKJ), Δ*htpG* (ΔG), and *ΔdnaKJΔhtpG* (ΔKJG) *E. coli* strains, the physiological role of HtpG and its possible collaboration with DnaK and the DnaJ co-chaperone, in unstressed *E. coli* cells grown at 30°C. Confirming earlier findings (Calloni, Chen et al. 2012), we observed that in the double ΔKJ mutant, few polypeptides were mildly, albeit significantly less soluble than in WT, but that mass-wise, an important population of DnaK and DnaJ substrates, including many metabolic and respiratory enzymes, was largely reduced, likely by proteases. Unexpectedly, the decreased cellular levels of these aggregation-prone polypeptides was significantly less pronounced in the triple ΔKJG mutant, whose growth was also correspondingly improved at 37°C, compared to the double ΔKJ mutant. The data suggest that in unstressed bacteria, HtpG regulates the proper folding by DnaK and DnaJ of many polypeptides. But when DnaK and DnaJ are inactive or transiently overwhelmed by misfolding polypeptides under heat stress, HtpG, which is strongly up-regulated, can promote an excessive and detrimental degradation of misfolding polypeptides, preferably by the HslUV protease. Because contrary to DnaK-DnaJ-GrpE-ClpB chaperones that can disaggregate and refold proteins with compromised structures (Goloubinoff, Mogk et al. 1999, Diamant, Ben-Zvi et al. 2000), HtpG-driven proteolysis by HslUV would be irreversible, this could affect the cellular function of essential proteins, limiting bacterial growth and increasing its sensitivity to heat stress.

## Results

### Mutation in htpG partially suppresses the growth defect of the double ΔdnaKdnaJ mutant at 37°C

In order to address a potential synergy between the DnaKJ chaperone machine and HtpG, we first designed a set of isogenic *E. coli* mutants in W3110 (WT) strain background, namely W3110 *ΔdnaKdnaJ* (ΔKJ) W3110 *ΔhtpG* (ΔG) and W3110 *ΔdnaKdnaJhtpG* (ΔKJG) and tested their ability to grow at various temperatures. No major differences in growth were observed at 30°C. At higher temperatures, ΔKJ growth was severely affected, as initially shown by Bukau and Walker (Bukau and Walker 1989) (**Figure 1, Figure S1**). Strikingly, we observed at 37°C that the triple ΔKJG mutant grew significantly better than the ΔKJ mutant (**Figure 1A**), suggesting that the presence of endogenous HtpG was deleterious in a background lacking DnaK and DnaJ chaperones (**Figure 1B**). This harmful effect was confirmed the overexpression of HtpG from a plasmid, which dramatically inhibited the growth of ΔKJ but not of WT, already at 30°C (**Figure 1C**). In contrast, the overexpression of HtpG affected less the growth of ΔKJG at 30°C (**Figure 1C**), possibly because it contained sevenfold less HtpG than the ΔKJ mutant (**Table S1**).

**Figure 1:**
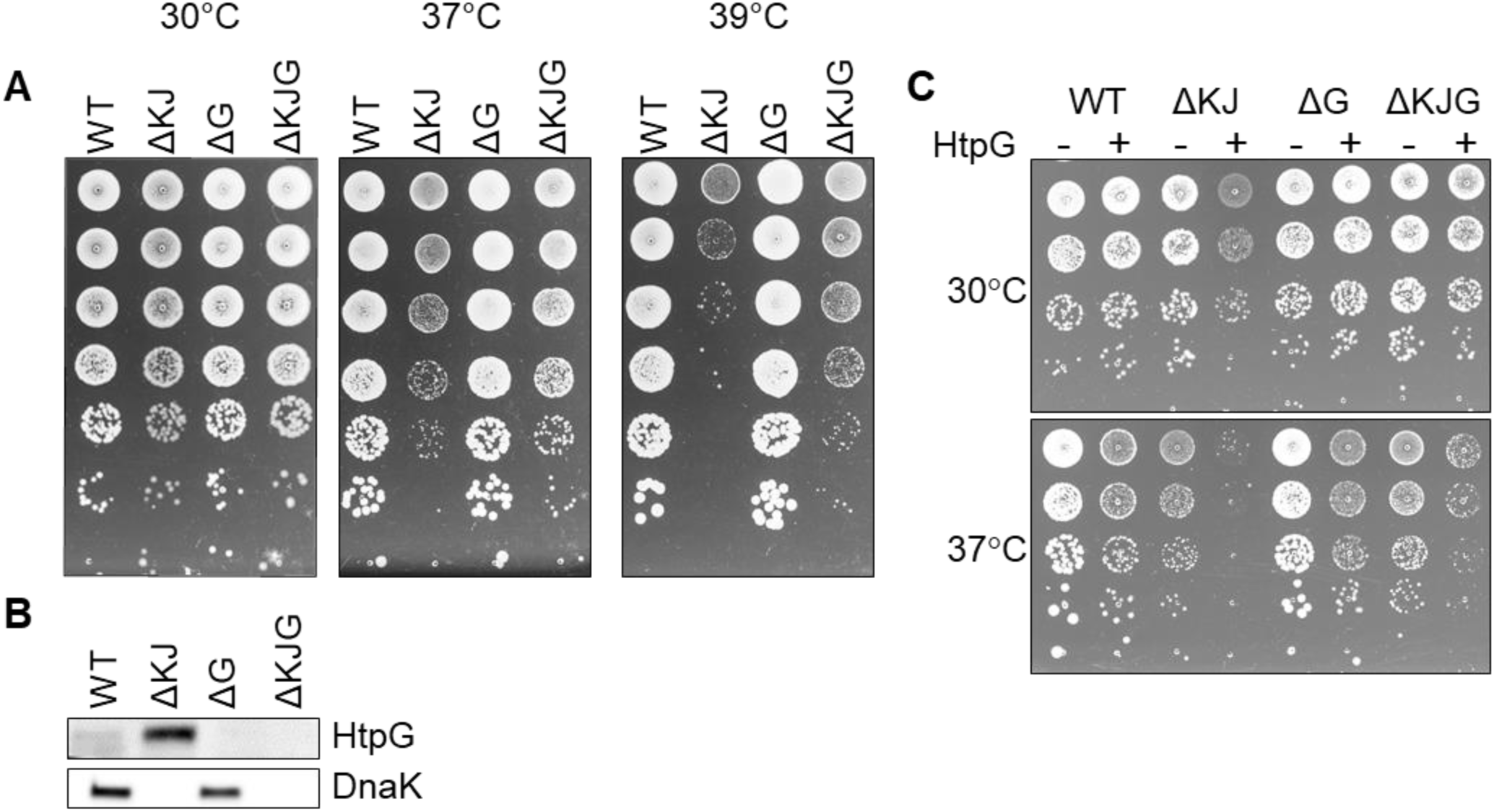
Mutation in htpG partially suppresses the growth defect of a ΔKJ mutant. **A:** Mid-log phase cultures of WT W3110 *E. coli* cells, ΔKJ, ΔG and ΔKJG *E. coli* strains were serially diluted 10-fold and spotted on LB agar plates and grown O/N at the indicated temperatures. **B:** Western blots of whole cell extracts using anti-DnaK and anti-HtpG antibodies. **C:** Toxicity of HtpG overexpression in ΔKJ background. Fresh transformants of W3110 derivative strains containing either an empty plasmid (pSE380-LacZ), or HtpG (pSE380-htpG) were grown at 30°C, serially diluted 10-fold and spotted on LB ampicillin agar plates supplemented with 1 mM IPTG for induction of HtpG overexpression. Plates were incubated for 1 day at 30°C (top panel) or 37°C (bottom panel).

We next used label-free MS-based absolute quantitative proteomic analysis, in an attempt to address and further identify particular *E. coli* polypeptides that specifically accumulate, degrade and/or aggregate in cells lacking, either HtpG, DnaK&DnaJ or all three. We identified and quantified the most abundant proteins in the total protein fractions of WT, ΔKJ, ΔG and ΔKJG strains grown at 30°C, and quantified their corresponding amounts in insoluble fractions. To optimise the statistical significance of the mean quantitative values obtained for each identified protein, we analysed by MS five independent biological samples of total (soluble and insoluble) proteins and the five corresponding insoluble-only fractions, for each of the four strains: WT, ΔKJ, ΔG and ΔKJG, a total of forty separately grown and independently prepared protein samples.

The total protein fractions from the five biological samples of WT *E. coli* cells grown separately at 30°C were first analyzed. Quantification was performed on the basis of iBAQ and LFQ intensities (Schwanhausser, Busse et al. 2011, Cox, Hein et al. 2014), which were first normalized into mass fractions *(i.e*. how much each protein contributes to the total protein mass per cell). We then took advantage of our five biological replicates to filter out proteins with very low average abundance and high variance *via t* tests with multiple testing correction (see materials and methods). Of the estimated ~4300 putative open reading frames of the *E. coli* genome (Kitagawa, Ara et al. 2005), 1203 proteins with significant mean mass fraction values were identified and quantified (their False Discovery Rate values were below 0.01, **Table S2**). Although being only a third of the total gene-encoding potential of the bacterial genome, these 1203 most abundant proteins summed up to be 97.2 % of the total protein mass of the cell. A scatter plot of the individual mass fractions from these proteins (**Figure 2A**) showed that the pellet mass fractions of the 281 *bona fide* identified membrane proteins (**Figure 2, orange dots**) were systematically higher than their corresponding total mass fractions: the sum of the total mass fractions of these 281 proteins represented 4.25 % of the total protein mass (**Table S2**); however, in the pellet fractions they summed up to 28 % of the WT *E. coli* pellet sample. This ~6.5-fold enrichment of pellet mass fractions compared to the true amounts in total cell lysates was confirmed and refined by a regression analysis of the total and pellet mass fractions of these membrane proteins. On the basis of the fitting performed on the 281 membrane proteins, all pellet mass fraction values were scaled down by a factor of ~6.5 for WT *E. coli*. The corrected scatter plot (**Figure 2B**) showed that the known membrane proteins then became positioned close to the 45° diagonal, which is the expected position of fully water-insoluble proteins, such as water-insoluble membrane-spanning and large insoluble structural proteins (**Figure 2, orange dots**). This procedure was then performed on the data from the other three *E. coli* strains, so as to perform meaningful comparisons between total and pellet mass fractions in each strain (**Table S3** and **Figure S3**). The corrected plot provided a relative solubility index for each significantly quantified protein. Because of the relatively low degree of significance of these solubility index values, and for simplicity, we defined here as <u>fully soluble</u>, all proteins that were found to be at least 90% soluble (**Figure 2, blue dots, below the hatched line**) and as the <u>fully insoluble</u>, all proteins that found in the plot to be less than 30% soluble (**Figure 2**, *gray and orange dots, above the hatched line)*.

**Figure 2:**
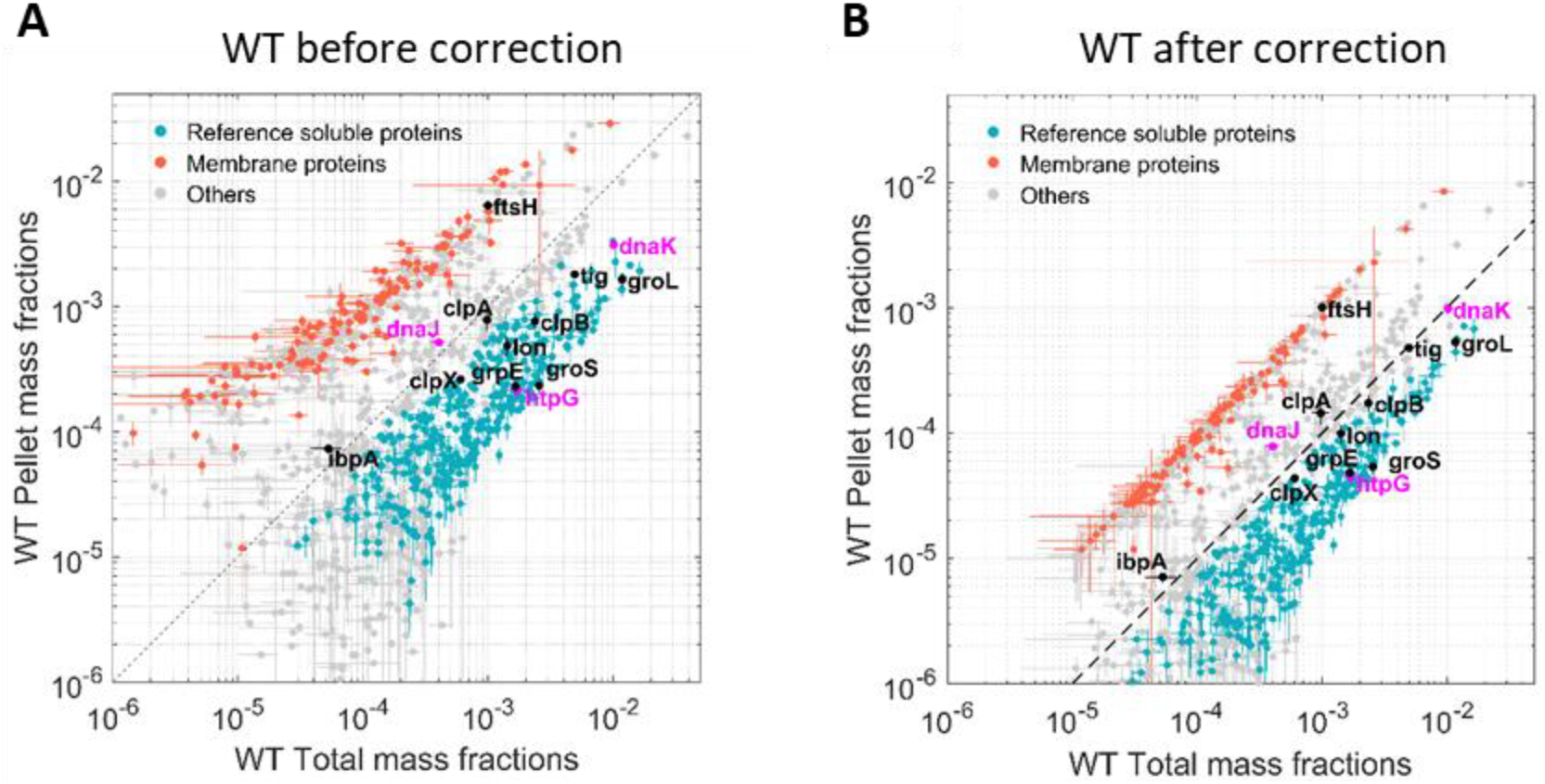
Quantitation of the most abundant soluble and insoluble proteins in WT *E. coli* cells. Scatter plot of the relative mean mass fractions of 1421 proteins from wild type W3110 *E. coli* that were significantly quantified in the total fraction containing both soluble and insoluble proteins (X-axis), against only in the insoluble pellet fraction, containing mostly the insoluble proteins (Y-axis:). **A:** Before correction for the over-representation of the insoluble proteins in the pellet fractions, **B:** After correction. Orange dots: 281 membrane proteins identified as such by the Uniprot database. Blue dots bellow the hatched line: proteins with a significant solubility index equal or greater than 90%. Grey circles: other less significantly soluble proteins. Magenta dots: DnaK, DnaJ and HtpG. Black dots, other canonical chaperones and major proteases.

To validate this calculation, we verified that DjlA, a membrane-anchored co-chaperone of DnaK and FtsH, a membrane-embedded essential protease, were both located among the insoluble membrane proteins. In contrast, most other classical intracellular molecular chaperones and proteases, which are acknowledged soluble proteins, were above the 90% solubility threshold (**Figure 2**, black and blue dots). Noticeably, IbpA and DnaJ appeared slightly less soluble (**Figure 2**, grey dots), but given that IbpA can assembles into variably large oligomers (Kitagawa, Miyakawa et al. 2002) and that both are known to associate with misfolded and insoluble aggregates, they might turn up being partly less soluble, even in WT cells grown at 30°C.

Taking an estimated total protein concentration in *E. coli* cells of 235 mg/mL (Zimmerman and Trach 1991, Ellis 2001) and knowing the specific molecular weight of each of the quantified polypeptide, we could translate the relative mean mass fraction values into cellular concentration estimates, expressed in μM (protomers). Thus, in WT cells grown at 30°C, the concentrations of DnaK, DnaJ and HtpG were, respectively, 32.6, 2.4 and 5.4 μM protomers. There were 46 μM GroEL protomers and 56 μM GroES protomers (**Table S1**), corresponding to about 3 μM of so-called footballs, which are functional GroEL1_4_[GroES_7_]2 oligomers capped on both sides by GroES_7_ (Azem, Diamant et al. 1995). Validating our methodological approach to convert LFQ MS data (Cox, Hein et al. 2014) into protein cellular concentrations, the concentrations deduced from the MS data showed a very good correlation with five previously published estimated concentrations of *E. coli* proteins obtained by similar and non-MS methods (Craig, Cortens et al. 2004, Valgepea, Adamberg et al. 2010, Arike, Valgepea et al. 2012, Krug, Carpy et al. 2013, Schmidt, Kochanowski et al. 2016) (**Figure S2**). We then applied the same statistical analysis to total and insoluble fractions of the ΔKJ, ΔG and ΔKJG strains, which produced, respectively, 1061, 1156, 1056 significantly quantified proteins, summing up to be, respectively, 96.7%, 97.3% and 96.7% of the total protein mass of the cells (**Figure S3**).

### Analysis of significant differences in protein concentrations between WT and mutant strains

Our data provided precise protein concentrations with high statistical significance for WT and the three mutant strains. Compared to WT, the ΔKJ deletion caused a massive net significant accumulation of 635 proteins accounting for 26.9 % of the total protein mass that was counterbalanced by net significant mass loss in 511 proteins, accounting for 26.7% of the total protein mass (**Table S2**). Our data confirmed earlier quantitative proteomic data on ΔKJ by Calloni et al. (Calloni, Chen et al. 2012), showing with SILAC-based MS analysis, that in the absence of DnaK and DnaJ, there was a dramatic increase in the cellular concentrations of the other remaining molecular chaperones and proteases. We found that the whole chaperone load of ΔKJ cells was increased from 3.2% to 8.8%, despite the loss of DnaK and DnaJ that accounted in WT cells for ~1% of the total protein mass (**Figure 3 A, B, Table S2**). A similar increase of the chaperone load was observed in the triple mutant ΔKJG, indicating that HtpG is not involved in the degradation of the σ^32^ transcription factor, at variance with DnaK and DnaJ (Lim, Miyazaki et al. 2013). A close-up analysis of the bacterial “chaperome” (**Figure 3B, Table S1**) showed that the complete deletion of *dnaK and dnaJ in* ΔKJ was counterbalanced by a three-fold increase in GroEL and GroES, and a 7-fold increase in HtpG and ClpB (**Table S1**). The small Hsps, IbpA and IbpB, which were virtually undetected in WT, were massively accumulated in ΔKJ (63- and ~2000-folds for IbpA and IbpB, respectively) and in ΔKJG mutants, but not in ΔG. Noticeably, large fold-change values did not necessarily translate into as extensive variations of protein abundances. Thus, although IbpB was upregulated ~2000-fold in ΔKJ, it merely contributed a net 0.2% increase to the total protein mass of the cell. In contrast, GroEL, which was upregulated merely 3.4-folds, contributed a considerable net 2.4 % increase to the total protein mass of the cell, i.e. 12 times more than IbpB.

**Figure 3:**
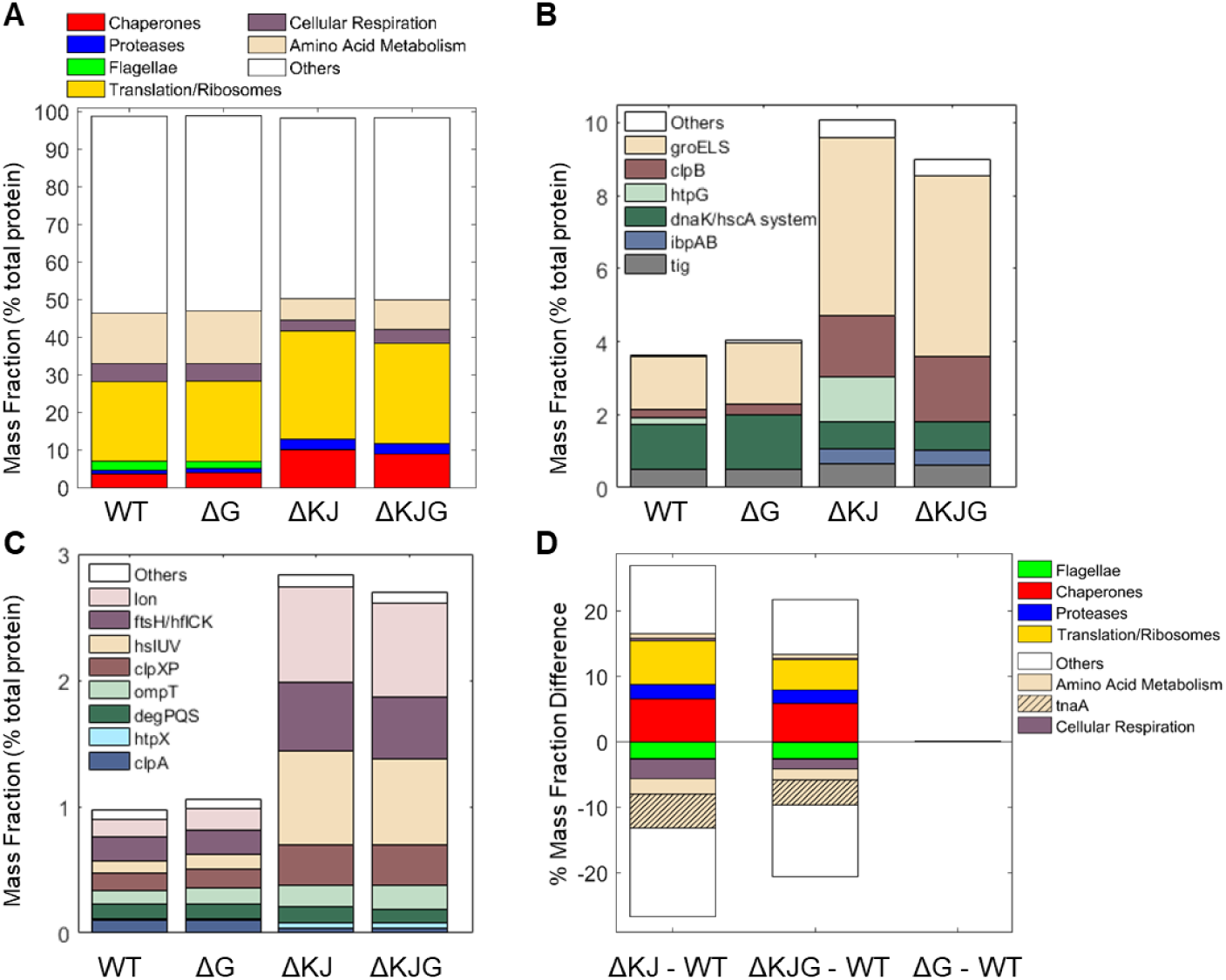
Analysis of different protein functional categories in the *E. coli* variants. **A:** Distribution of the principal functional categories of significantly quantified proteins in the four *E. coli* strains, Data are presented as mass fractions. Functional categories were manually defined according to both previous knowledge from the literature and GO annotations. **B:** Mass fractions of individual proteins belonging to the chaperone category. The dnaK/hscA system (dark green bar) comprises dnaK, dnaJ, grpE, cbpA, djlA, hscA, and hscB. The white bar (‘Others’) comprises hslO, and spy. **C:** Mass fractions of individual proteins belonging to the protease category. **D:** Mass fraction differences (expressed in % of the total intracellular proteins) between WT and the various deletion mutants. When absent in their respective mutants, the proteins DnaK, DnaJ, and HtpG were excluded from the analysis.

Similar to classical chaperones, but mass-wise to a lesser extent, ATP-dependent proteases, such as Lon, FtsH, ClpXP and HslUV also accumulated between 3 to 9-fold in ΔKJ and ΔKJG, but not in ΔG (**Figure 3C; Table S1**). Only ClpA, which like ClpX associates to the ClpP protease, was less expressed in ΔKJ, in agreement with ClpA expression not being under the control of σ^32^ (Katayama, Gottesman et al. 1988) and with ClpXP, Lon, FtsH, and/or HslUV likely being involved in the degradation of proteins who failed to properly fold in the ΔKJ mutants. Noticeably, as previously shown by Calloni and colleagues (Calloni, Chen et al. 2012), DegPQS, which are non-ATPase periplasmic proteases and therefore are not expected to be involved in the degradation of cytosolic proteins, were less expressed in ΔKJ than in WT.

Both in ΔKJ and ΔKJG strains, there was also a marked increase of ribosomal proteins and of associated components of the protein translation machinery, as compared to WT: their proportion massively increased from 19.9% of the total mass in WT, to 27.2% in ΔKJ and to 25.1% in ΔKJG. Because in ΔKJ, the amount of insoluble proteins at 30°C was not markedly higher than in WT, this implies that the mass gain from the newly synthetized chaperones, proteases and ribosomes, must have been counterbalanced by a corresponding mass loss from the synthesis-arrest and/or from the specific degradation by proteases of other abundant proteins, such as enzymes from the amino acid metabolism. This is in agreement with the major role that was initially found for DnaK in central metabolism (Angles, Castanie-Cornet et al. 2017) by microarray work (Fan, Liu et al. 2016), and with the earlier SILAC MS proteomic observations that many DnaK-DnaJ-dependent substrates become significantly degraded in ΔKJ, as compared to WT cells (Calloni, Chen et al. 2012).

We found that the remarkable high levels of metabolic enzyme tryptophanase (TnaA) in WT were strongly decreased from 6.4% to 0.9 % of the total protein mass in ΔKJ. Other abundant proteins, such as the known DnaK interactors PflB and PutA (Calloni et al, 2012) (**Table S2**) were also decreased in ΔKJ. In addition, confirming early reports that flagellum synthesis strictly depends on DnaK (Shi, Zhou et al. 1992), we found that several members of flagellar machinery proteins, principally flagellin (FliC) were completely absent in ΔKJ. Interestingly, already at this coarse level of analysis of customized and GO protein categories (**Figure 3A**), we observed almost no differences in the mass profile of the proteins in ΔG, as compared to the WT strain. The only exception was ClpB, which out of ~1300 significantly quantified proteins in both WT and ΔG strains, was mildly more abundant in ΔG. The lack of proteomic phenotype in unstressed ΔG cells was confirmed by the triple mutant ΔKJG, which showed a protein profile quite similar to that of the double ΔKJ mutant. Yet, the extent of mass loss in enzymes involved in amino acid metabolism and in cellular respiration was found to be systematically less pronounced in ΔKJG (**Figure 3D**). This suggests that there is an inverse correlation between growth and the relative mass loss of key metabolic enzymes (Angles, Castanie-Cornet et al. 2017) in the ΔKJ mutant, as compared to ΔKJG.

The observation by mass spectrometry that the endogenous level of TnaA was severely reduced in ΔKJ, compared to WT and that it was less reduced in ΔKJG, was confirmed by Western Blot (WB) analysis of total cell lysates of the four W3110 *E. coli* variants. When cells were grown at 30°C and sampled at mid-log phase, precisely as in the proteomic analyses, the TnaA abundance was found to be reduced in ΔKJ to 55 ± 11% of the WT levels (**Figure 4 A, B**). The observed loss of TnaA in ΔKJ was more pronounced in cells cultured at 37°C: WB analysis showed a TnaA level of 19 ± 8 % (relative to WT) in ΔKJ (**Figure 4 A, B**) This result indicates that in exponential phase of growth, TnaA cannot reach its native state without assistance by DnaK and DnaJ and instead becomes targeted by the ~8 fold higher levels of HtpG to degradation by proteases, which are more than twice more abundant. Our observation by MS of a partial recovery of TnaA levels in ΔKJG (37 ± 7 % of WT) compared to ΔKJ (13 ± 3 % of WT) was not as clear by western blot, where at 30°C TnaA levels were 50.2 ± 9.6% of WT and 55.4 ± 11.2 % of WT in ΔKJG and ΔKJ, respectively; and at 37°C TnaA was 18.8 ± 17.5% of WT and 19 ± 8 % of WT in ΔKJG and ΔKJ, respectively (**Figure 4B**). This could be due to the lower accuracy of the western blot semi-quantitation, compared to MS.

**Figure 4:**
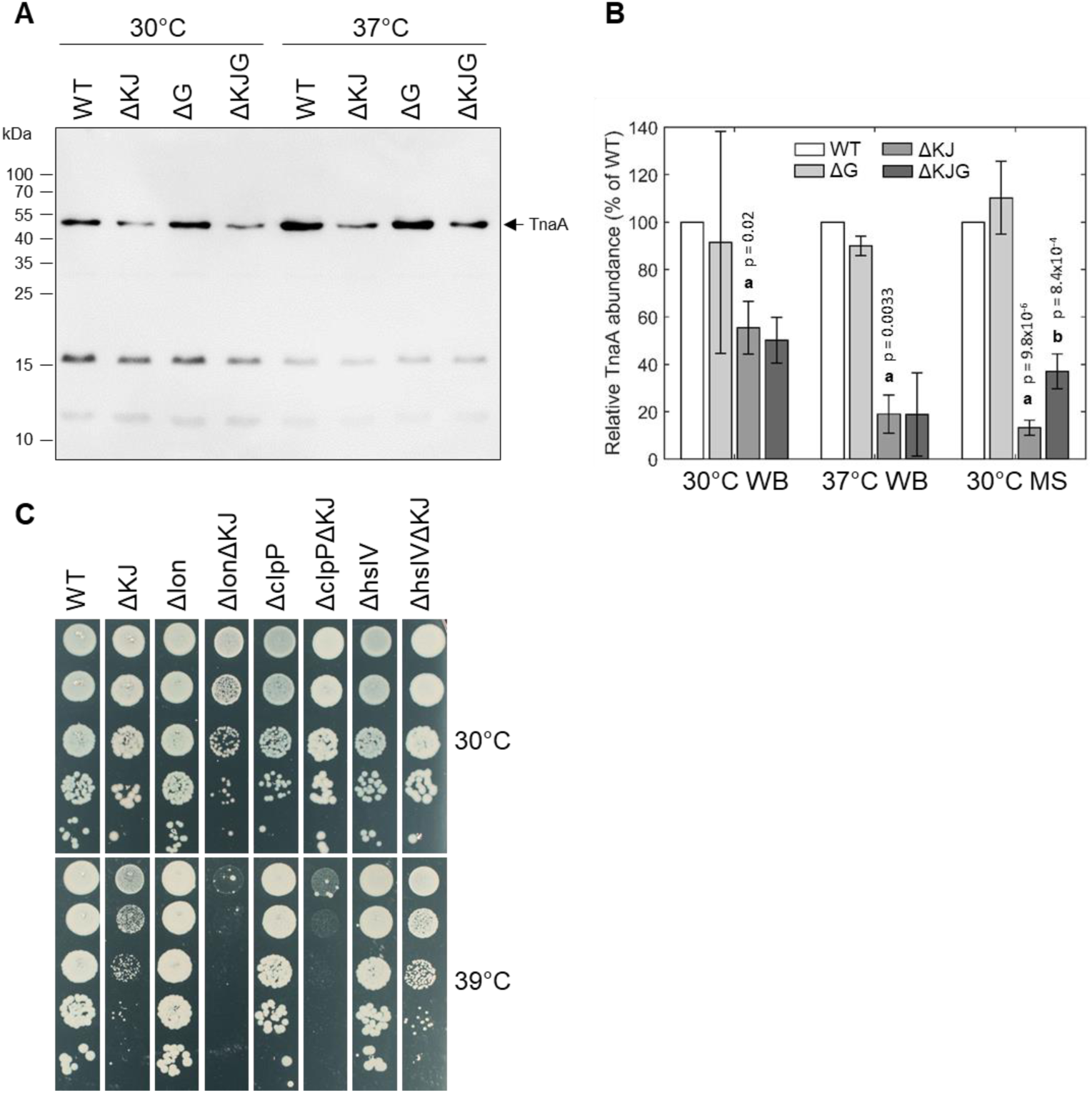
Validation of TnaA degradation by Western Blot. **A:** Representative semiquantitative SDS-PAGE/Western Blot of W3110 *E. coli* lysates. **B:** Densitometry analysis of the Western Blot shown in A. A total of n=3 biological replicates were performed for Western Blot analysis. The dark gray bars correspond to the relative (compared to WT) tryptophanase abundances as determined by proteomics analysis (see Fig. 5). Letters above the bars indicate significant statistical comparisons. a: Significant difference between WT and ΔKJ. b: Significant difference between ΔKJ and ΔKJG. p-values are indicated above. **C:** hslV deletion improves growth in the ΔKJ background. Serially 10-fold diluted cultures of WT, ΔKJ, or ΔKJ combined with the indicated single-protease deletions (Δlon, ΔclpP and ΔhslV) were spotted on agar plates and grown overnight at 30°C (top) or 39°C (bottom).

We next addressed genetically, which of the main *E. coli* proteases might be involved in the HtpG-promoted degradation of misfolded DnaKJ substrates. The triple ΔKJ-protease mutants *ΔdnaKdnaJΔlon, ΔdnaKdnaJΔhslV*, and *ΔdnaKdnaJΔclpP* were grown at different temperatures. At 30°C, the deletion *of lon* impaired the growth of the ΔKJ strain (**Figure 4C**). At 39°C, the deletion *of lon* or *clpP* further impaired ΔKJ growth, whereas, remarkably, deletion of *hslV* partially suppressed the growth defect of the ΔKJ strain (**Figure 4C**). Together with the results from the pulse-chase SILAC MS experiments from Calloni *et al*. (Calloni, Chen et al. 2012), who showed increased protein degradations in ΔKJ compared to WT, and the TnaA western blots (**Figure 4A, B**), our finding that HslV deletion improved growth of ΔKJ suggests that HtpG is involved in mediating the degradation, principally *via* the HslUV protease, of aggregation-prone polypeptide that need DnaK-DnaJ to properly fold. The 935 proteins that significantly differed in ΔKJ from WT and the 834 proteins that significantly differed in ΔKJG from WT were next sorted according to protein categories. In both groups, chaperones, proteases and proteins of the translation machinery were almost exclusively upregulated, whereas flagellar proteins, amino acid metabolic enzymes, and proteins of the energy metabolism (cellular respiration) were mostly down-regulated (or degraded).

It was initially shown that compared to WT cells, ΔK cells massively accumulate insoluble proteins at 42°C, but not at 30°C (Mogk, Tomoyasu et al. 1999). Here, we found that in WT, ΔKJ and ΔKJG strains grown at 30°C, the sum of the insoluble proteins was, respectively, 18.5%, 20.5% and 18.9% of the total cellular proteins, suggesting that at 30°C the ΔKJ background contained slightly but more dramatically more insoluble aggregates than the WT. Solubility differences were then assessed in a more stringent list obtained with *t* tests (test with multiple testing correction, FDR cutoff 0.05) to determine which proteins were statistically differing in their solubility between WT and ΔKJ. The list was furthermore restricted to proteins with more than 10 percent point difference in solubility *(i.e*. | Solubility_WT_ - Solubility_Δ_KJ | > 0.1 [where Solubility = (1 - Pellet)/Total], because smaller solubility differences are unlikely to hold biological significance). We thus obtained a list of 161 proteins (**Table S4**), 133 of which were less soluble in ΔKJ than in WT. These 133 proteins, which summed up to 7.3%, 10.4%, and 9.8% of the total protein mass in WT, ΔKJ, and ΔKJG, respectively. Their average solubility decreased from 91% in WT to 70% in ΔKJ. The solubility of these proteins did not change in ΔKJG, being on average 71% as in ΔKJ.

The proteomic analysis of the ΔKJ mutant thus indicated that, expectedly, ΔKJ contained somewhat less soluble proteins, although in relatively small numbers and contribution to the total proteome, which is consistent with the observation that at 30°C, the ΔKJ cells grew similarly to WT cells.

Chaperones are classically described as molecules that prevent protein aggregation. We therefore next compared the effects of the HtpG over-expression, or its deletion into the ΔKJ background, on the specific solubility indexes of key individual proteins and their total cellular amounts. The 40 most abundant proteins that most significantly accumulated in ΔKJ compared to WT were either ribosomal proteins, chaperones (**Figure 5B**, *red arrowheads*) or proteases (**Figure 5B**, *blue arrowheads*). Whereas, together they summed up to be 18.3% of the total protein mass of WT cells, they were nearly doubled (33.4%) in ΔKJ cells and slightly less (31.6%) in ΔKJG cells. Remarkably, although they were significantly less abundant in WT and ΔKJG than in ΔKJ, they all maintained the same degree of relative solubility, **(Figure 5B, D, F**). Thus, the ΔKJ phenotype cannot be attributed to a decrease in the solubility of the proteins that became most up-regulated in ΔKJ.

**Figure 5:**
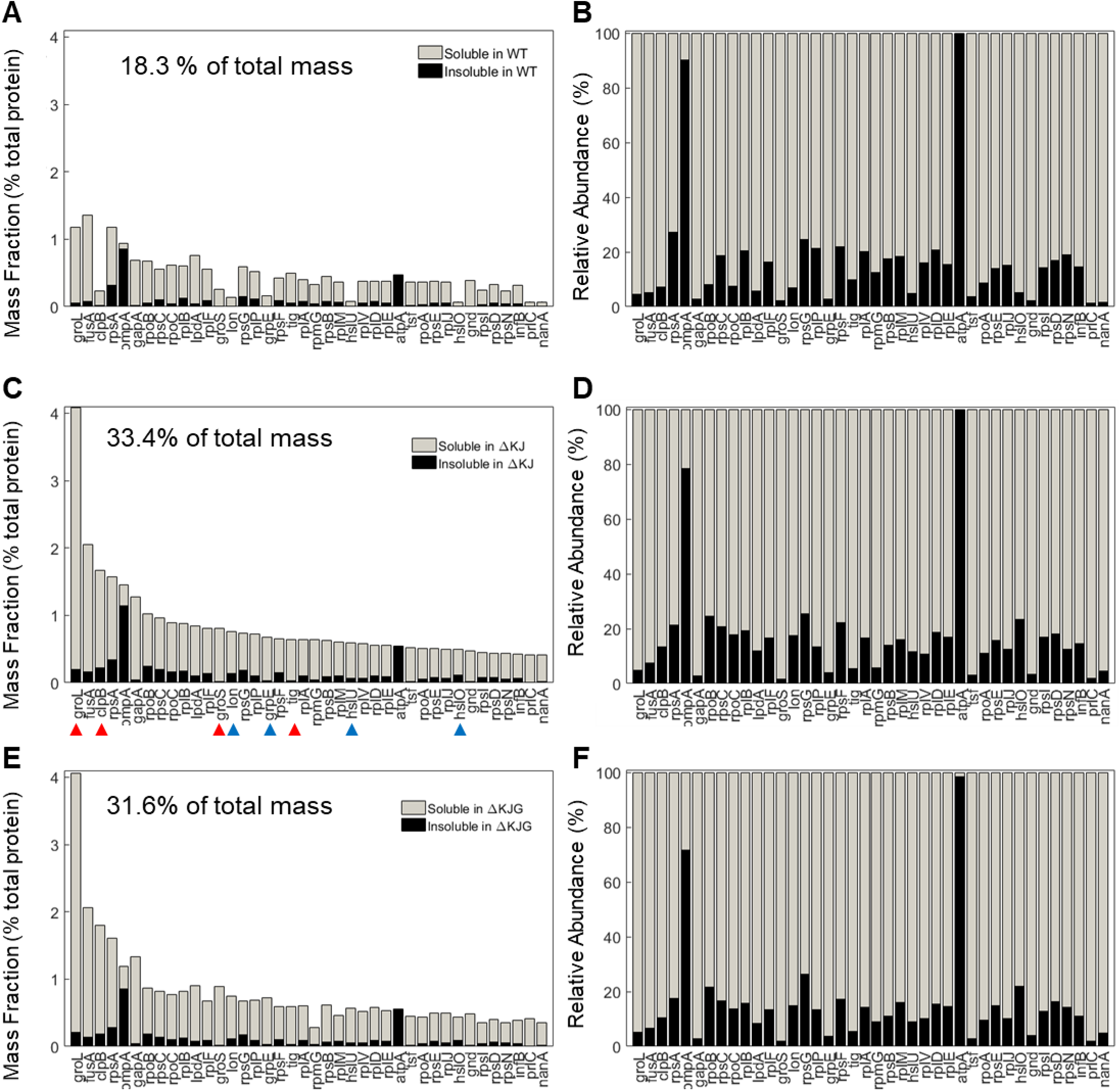
Mild aggregation of the most abundant proteins significantly upregulated in ΔKJ. **A:** Absolute abundances (mass fractions of the total cellular protein amount) in WT soluble (grey bars) and insoluble (black bars) fractions for the 40 most abundant proteins of ΔKJ *E. coli* that were significantly upregulated in ΔKJ compared to WT. **B:** Normalized abundances in WT E.coli of the same proteins as in A. **C:** Absolute abundances (mass fractions) of the same proteins as in A, but showing their amounts in ΔKJ soluble (grey bars) and insoluble (black bars) fractions **D:** Same as in C, with normalized abundances. E: Absolute abundances (mass fractions) of the same proteins as in A, but showing their amounts in ΔKJG soluble (grey bars) and insoluble (black bars) fractions. **F:** Same as in E, with normalized abundances.

The involvement of molecular chaperones, DnaK in particular, in intracellular protein degradation of proteins has long been documented (Sherman and Goldberg 1996) and recently confirmed in greater details by SILAC-MS with pulse-chase experiments (Calloni, Chen et al. 2012). We therefore next identified the 40 most abundant proteins in WT (**Figure 6 A, B**) that were significantly less abundant in ΔKJ (**Figure 6 C, D**) and found that most were enzymes involved in cellular respiration and in the amino acid metabolism, in particular tryptophanase A (TnaA). Together, the mass of these 40 most reduced proteins in ΔKJ compared to WT, was 26.8% in WT, 9.1% in ΔKJ and 13.7% in ΔKJG, corresponding to a dramatic three-fold mass loss in ΔKJ, compared to WT, which was lessened to a two-fold in ΔKJG. Indeed, 39 of these 40 proteins were found to be significantly more abundant in ΔKJG (**Figure 6 E, F**) than in ΔKJ (**Figure 6 C, D**), further confirming that the deletion of htpG leads to less degradation of these proteins that otherwise tend to misfold in the ΔKJ background.

**Figure 6:**
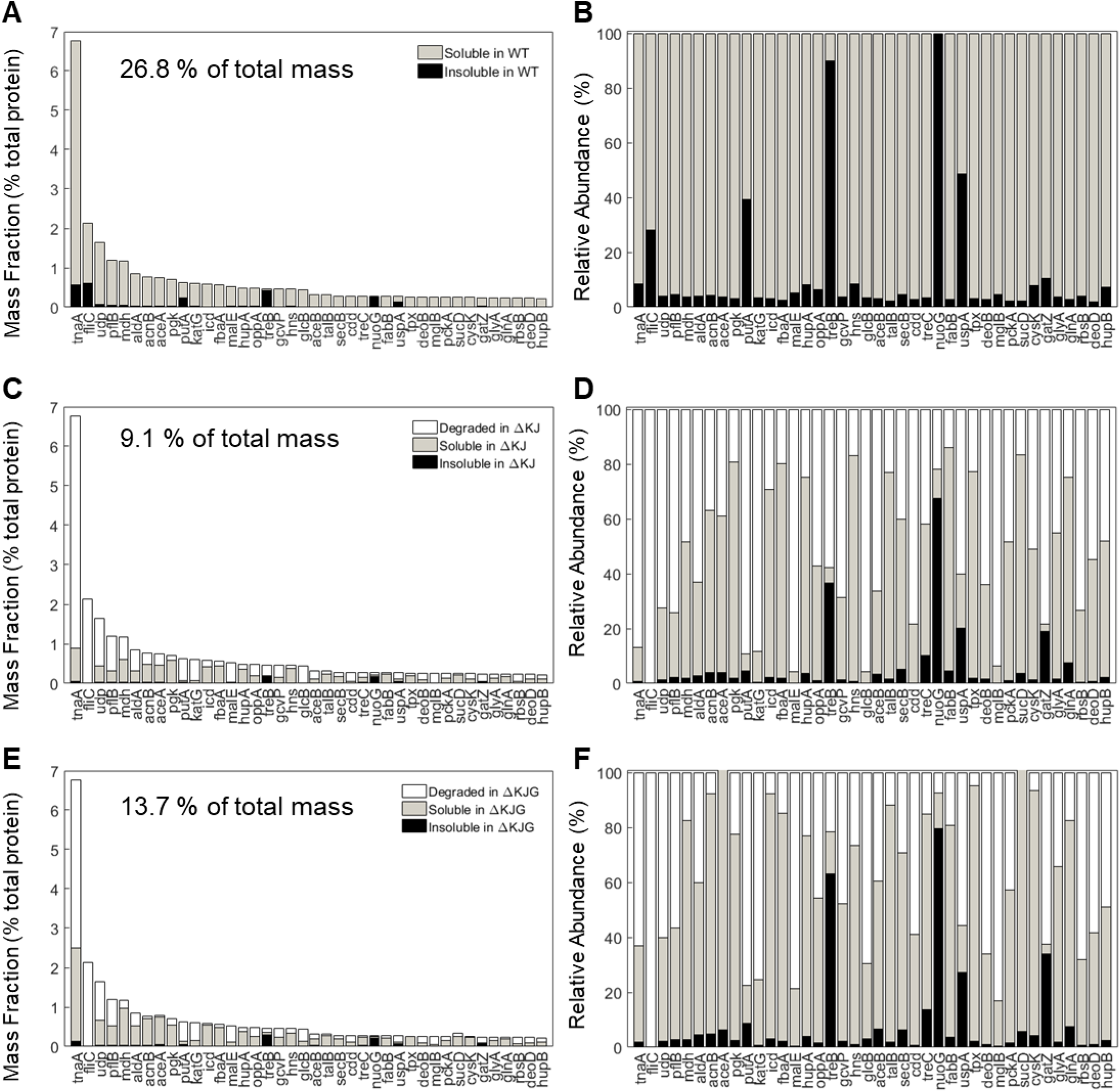
Protein degradation is a prevalent phenotype in ΔKJ and partially rescued in ΔKJG. **A:** Absolute abundances (mass fractions of the total cellular protein amount) in WT soluble and insoluble fractions for the 40 most abundant proteins of WT *E. coli* that were downregulated in ΔKJ. **B:** Normalized abundances in WT *E. coli* of the same proteins as in A. **C:** Absolute abundances of the same proteins as in A, but with a partitioning showing the amounts of each protein degraded in ΔKJ (white bars); the soluble (grey bars) and insoluble (black bars) amounts in. **D:** Same as in C, with normalized abundances. **E:** Same absolute abundances as in A and C, but partitioned in order to show the quantities degraded in ΔKJG with respect to WT (white bars); the soluble (grey bars) and insoluble (black bars) amounts in ΔKJG. **F:** Same as in E, with normalized abundances.

## Discussion

The Hsp90s are members of a highly conserved family of ATP-hydrolyzing molecular chaperones, which together with the Hsp70s form the core of the cellular protein quality control machinery (Taipale, Jarosz et al. 2010, Balchin, Hayer-Hartl et al. 2016). The various cytosolic and organellar orthologues of Hsp90 are among to most abundant proteins of animal cells, reaching ~ 1 % of the total protein mass (Finka and Goloubinoff 2013, Gat-Yablonski, Finka et al. 2016). Here, we find that in bacteria, DnaK levels are also ~ 1 % as in the cytosol but that HtpG levels are generally 7 times lower, unless DnaK is deleted (Figure S4).

Little is known about the precise molecular mechanism by which Hsp90s carry their specific ATP-fuelled protein-restructuring functions. It is not known why are they so evolutionarily conserved, and what makes them particularly essential under both physiological and stress conditions, compared to other types of molecular chaperones. Similar to Hsp60s (GroEL), Hsp70s (DnaK), Hsp40s (DnaJ) and small Hsps (IbpA, IbpB), Hsp90s (HtpG) (E. coli chaperones in brackets) can bind misfolding polypeptides and prevent the formation of inactive and potentially toxic protein aggregates (Stefani and Dobson 2003). Whereas Hsp60/10, Hsp70/40, Hsp104/70 are unfolding enzymes that can use the energy of ATP hydrolysis to forcefully translocate polypeptide to convert stable misfolded or oligomeric protein complexes into native proteins, very little is known about the particular nature of the structural changes applied by the Hsp90s onto their bound protein clients (Finka, Mattoo et al. 2016). Hsp90 assembles into very dynamic dimers (Shiau, Harris et al. 2006, Krukenberg, Street et al. 2011, Li, Soroka et al. 2012, Flynn, Mishra et al. 2015, Mayer and Le Breton 2015). There are profound structural differences between the apo-, ATP-and ADP-bound states. A central question about the chaperone mechanism of Hsp90s is the role of specific co-chaperones. In the cytosol of eukaryotes, Hop, Aha1 and P23 are three conserved Hsp90 co-chaperones controlling precise structural changes in the Hsp90 dimer during the various stages of the ATPase cycle (Mayer and Le Breton 2015). Yet, there are no apparent orthologues of Hop, Aha1 and P23 in the lumen of the endoplasmic reticulum, in the stroma of mitochondria or in the cytosol of eubacteria, to co-chaperone, respectively, GRP94, Trap1 and HtpG. In contrast, Hsp70 is always co-expressed with Hsp90s in all the ATP-containing compartments of cells, raising the possibility that Hsp70 is acting as the most conserved co-chaperone to all prokaryotic, organellar and eukaryotic forms of Hsp90s (Finka, Mattoo et al. 2011). There are numerous examples of Hsp90 and Hsp70 acting together, as in the case of steroid hormone receptor activation (Echeverria and Picard 2010) and in a context devoid of a Hop cochaperone, as in *E. coli* cells (Nakamoto, Fujita et al. 2014). *In vitro*, HtpG from *E. coli* and cyanobacteria seem to collaborate with the DnaK chaperone system to remodel denatured proteins (Genest, Hoskins et al. 2011, Nakamoto, Fujita et al. 2014). Whereas, much owing to the early finding of specific inhibitors, the involvement of Hsp90s in cellular signaling and stress protection has been demonstrated in eukaryotes, very little is known about the role of Hsp90s in proteobacteria, where it is apparently not essential.

In *E. coli*, deletion of the *htpG* gene, although highly conserved, leads only to a very minor phenotype observable only under elevated temperatures (Bardwell and Craig 1988, Thomas and Baneyx 2000, Grudniak, Pawlak et al. 2013, Press, Li et al. 2013). In cyanobacteria, it is involved in phycobilisome assembly (Motojima-Miyazaki, Yoshida et al. 2010, Sato, Minagawa et al. 2010, Press, Li et al. 2013), in heat stress (Thomas and Baneyx 1998, Thomas and Baneyx 2000) and it participates in oxidative stress resistance (Tanaka and Nakamoto 1999, Hossain and Nakamoto 2003). In pathogenic eubacteria, HtpG may be involved in bacterial immunity (Yosef, Goren et al. 2011) and virulence associated to toxin synthesis (Vivien, Megessier et al. 2005, Dang, Hu et al. 2011, Verbrugghe, Van Parys et al. 2015, Garcie, Tronnet et al. 2016). Honoré et al., recently found that HtpG from *S. oneidensis* is essential for growth at high temperature and identified tRNA-ileu synthetase as an essential HtpG client. Recent studies highlighted some functions of HtpG under stress conditions, where it is involved in stabilizing the essential protein TilS in the bacterium *Shewanella oneidensis* under heat stress (Honore, Mejean et al. 2017); as well as biofilm formation and secretion of certain enzymes such as β-lactamase in *E. coli* (Grudniak, Pawlak et al. 2013). Moreover, a recent genome-wide co-evolution study in *E. coli* predicted that under physiological conditions, HtpG may assist the folding of chemotaxis-related proteins, such as the chemoreceptor kinase CheA. HtpG was reported to be essential only in particular cases, such as for the activity of the CRISPR/Cas system (Yosef, Goren et al. 2011). Yet, the precise mechanism of action of the bacterial Hsp90, the identity of its substrates and its role under physiological and stress conditions remains elusive.

Here, we found that in unstressed *E. coli* cells at 30°C, the amount of HtpG was in general less abundant than in animal cells, only 0.15% of the total proteins, but that in ΔKJ cells, it was increased to 1%. Rationalising their traditional, albeit inaccurate, designation as heat-shock proteins (Hsps), this amount may increase by ~10% following a heat-shock, such as a fever (Finka, Sood et al. 2015) (Finka, Mattoo et al. 2011). Both Hsp90 and Hsp70 molecules can passively bind limited amounts of stress-labile polypeptides, thereby preventing some aggregations, albeit only to the limited extent of the individual binding (or so called “holdase”) capacity of each chaperone (Wiech, Buchner et al. 1992, Street, Zeng et al. 2014, Kravats, Doyle et al. 2017). Importantly, in the presence of an excess of misfolding polypeptides, Hsp70, Hsp104 and possibly Hsp90 too, can use the energy of ATP hydrolysis to unfold stably misfolded polypeptides and catalytically convert them into native proteins, even under stressful conditions that disfavour the native state (Genest, Hoskins et al. 2015, Finka, Mattoo et al. 2016, Goloubinoff, Sassi et al. 2018). As in the case of specific Hsp70s, elevated cellular levels Hsp90 are hallmarks of various types of stresses (Finka and Goloubinoff 2013, Finka, Sood et al. 2015). At 41°C, human cell cultures can accumulate 50% more Hsp90s, compared to 37°C (Finka, Sood et al. 2015). Likewise, when naive rat liver cells at 37°C are being stressed by chronic excessive food intake (**Figure S4**, AL), they constitutively express ~50% more Hsp90s and Hsp70s than unstressed cells from mildly food-restricted livers (Gat-Yablonski, Finka et al. 2016). Justifying that Hsp90s and Hsp70s should be major drug targets for cancer therapy, various immortalized cancer lines constitutively over-express Hsp90s and Hsp70s at 37°C. Together, they can prevent the oligomerization and the activation of various client proteins, such as HSF1 (Abravaya, Myers et al. 1992), IΔB (Weiss, Bromberg et al. 2007) and block caspase activation (Multhoff 1997, Jaattela, Wissing et al. 1998, Lanneau, de Thonel et al. 2007, Neckers, Kern et al. 2007, Vartholomaiou, Echeverria et al. 2016), thereby promoting immortality of cancer cells and the general resistance to physical stresses and chemotherapeutic agents.

Here, we used an unbiased high-throughput quantitative proteomic approach of WT, and of DnaKJ and/or HtpG-knockout *E. coli* strains, ΔG, ΔKJ and ΔKJG, in an attempt to further understand the fundamental role of bacterial Hsp90, with its collaborating chaperones, Hsp70 and Hsp40, in a context devoid of the specific eukaryotic co-chaperones. We found that at non-stressful growth condition (30°C), bacterial Hsp90 was involved in the sorting of aggregation-prone proteins towards native refolding by the Hsp70/Hsp40 system, or in case of Hsp70/Hsp40 failure, towards more degradation preferably by the HslUV protease, as also shown by pulse-chase SILAC-MS experiments (Calloni, Chen et al. 2012). Accordingly, it has been demonstrated that in α-proteobacterium *Caulobacter crescentus*, DnaK depletion is suppressed by HslUV by attenuating σ^32^ factor (Schramm, Heinrich et al. 2017).

Our data may thus suggest a role for HtpG in the bacterial proteostasis network: it could identify proteins that fail to properly fold when DnaK is absent or overwhelmed by stress-induced aggregations. Even without a stress, whereas some nascent polypeptides can fold spontaneously to the native state, others are known to necessitate trigger factor (Tig) (Deuerling and Bukau 2004) (**Figure S5**). Others that misfold, for example in Tig mutants may need to be actively unfolded by bacterial Hsp70s and Hsp40s, in order for them to re-engage onto the native folding pathway to the functional state. Failing that in the ΔKJ knockout strain, where Tig is not dramatically more abundant, but where HtpG is seven times more abundant, misfolding proteins are redirected by HtpG to be degraded preferably by HslV, HslV and HslU being respectively 9 and 7.5 times more abundant in ΔKJ than in the wild type (**Table S1**).

A protein that has been degraded cannot be solubilized and reactivated by molecular chaperones. It must therefore be re-synthetized at an estimated ATP cost 100-1000 times higher than the cost of chaperone-mediated protein disaggregation and refolding (Sharma, De Los Rios et al. 2011). The impaired growth of ΔKJ at 37°C could thus be caused by to an excessive degradation of essential metabolic and respiratory proteins. Confirming that, the deletion of the *htpG* gene in ΔKJG (or of the *hslV* gene in ΔKJ), reduced protein degradation and restored bacterial growth at 37°C. This was further confirmed upon over-expressing HtpG in a ΔKJ background, which severely impaired ΔKJ growth, but not WT growth at 30°C.

The artificial over-expression of Hsp90 in *C. elegans* embryos was recently shown to cause the massive degradation of muscle proteins (Bar-Lavan, Shemesh et al. 2016), indicating that in the cytosol of stressed eukaryotic cells, Hsp90 might also mediate the specific degradation of misfolding proteins that escaped the action of other members of the chaperone network. It will now be interesting to address which are the particular protein partners in bacteria that mediate the interaction of Hsp90 and Hsp70 clients with the cellular proteases and further address the possibility that also in eukaryotes, Hsp90s could mediate the degradation by proteasome of Hsp70-clients that failed proper folding under stress or owing to mutations.

## Acknowledgements

This project was financed in part by the University of Lausanne and by the Swiss National Science Foundation Grants 140512/1, 31003A_156948 and by Grant C15.0042 from the Swiss State Secretariat for Education Research and Innovation.

## Material and Methods

### Bacterial Strains, phages, and culture conditions

Genetic experiments were carried out in the *E. coli* in W3110 genetic background strain (Bachmann 1972). The W3110 mutant derivatives *ΔdnaKdnaJ*∷Kan^R^; Δ*lon:*:Kan^R^ (Sakr et al., 2010) have been previously described. The *ΔhtpG:*:Kan^R^, Δ *lon:*:Kan^R^, Δ*clpP*∷Kan^R^ and *ΔhslV:*:Kan^R^ alleles were obtained from strains JWK0462, JWK0429, JWK0428, JWK0427 and JWK3903 (Keio collection). All mutations described in this study were moved to the appropriate genetic background by bacteriophage P1-mediated transduction. Bacteria were routinely grown in LB medium supplemented when necessary with either kanamycin (50 Δg/mL) or ampicillin (100 Δg/mL).

The construction of the *dnaK-protease* and *dnaK htpG* double mutants was performed as follows. The Kan^R^ cassettes from W3110 *ΔhtpG*∷Kan^R^, Δ *lon*∷Kan^R^, Δ*clpP*∷Kan^R^ and *ΔhslV*∷Kan^R^ were first removed using plasmid pCP20 as described (Datsenko and Wanner, 2000). The *ΔdnaKdnaJ*∷Kan^R^ mutant allele was then introduced into the *htpG* and the various protease mutants, thus leading to strains W3110 *ΔhtpG ΔdnaKdnaJ*∷Kan^R^, Δ*lon ΔdnaKdnaJ*∷Kan^R^, Δ*clpP* Δ*dnaKdnaJ*∷Kan^R^ and *ΔhslV ΔdnaKdnaJ*∷Kan^R^.

### Plasmid construction

Plasmid pSE380Δ *Nco*I (Genevaux, Keppel et al. 2004) has been previously described. To construct the high-copynumber plasmid pSE-HtpG (pSE380 Δ *Nco*I-HtpG), the 1875 bp *htpG* gene was PCR-amplified using primers HtpG-for(5′-CGGAATTCATGAAAGGACAAGAAACTCG-3′) and HtpG-rev (5′-CGAAGCTTTCAGGAAACCAGCAGCTGG-3′) using MG1655 genomic DNA as template. The PCR fragment was digested with *EcoRI* and *Hind*III and cloned into pSE380Δ *Nco*I previously digested with the same enzymes.

### Bacterial viability assays

Cultures of W3110 derivative strains were first grown overnight in LB at the permissive temperature (30°C for *ΔdnaKdnaJ*∷Kan^R^ strains and 37°C for the other strains), diluted 1/50 into the same medium, further grown to mid-log phase, serially diluted ten-fold and spotted on LB agar plates and incubated at the indicated temperatures. To monitor HtpG toxicity of the W3110 and its mutant derivatives, mid-log phase cultures of fresh transformants were grown at 30°C in LB ampicillin glucose 0.4%, were serially diluted ten-fold and spotted on LB ampicillin agar plates with or without IPTG (1 mM), and incubated at 30°C.

### TnaA SDS-PAGE / Western Blots

Cells were grown in 10 mL LB medium at 30°C until late stationary phase (overnight) or until mid-log-phase (OD ~0.4), and then centrifuged at 7000g for 10 min at 4°C. Cell pellets were resuspended in 400 μL lysis buffer (50 mM Tris pH 8.0, 150 mM NaCl, 1 mM PMSF, and 200-times diluted protease inhibitor cocktail (Sigma, cat. # P8465) and lysed on ice by sonication (2 cycles, each with 30 pulses at 80% amplitude and 50% duty cycle). Total concentrations were then normalized with a Bradford assay before loading 5 μg total protein for each sample on 12% polyacrylamide SDS-PAGE gels. Following migration (160 V), proteins were transferred onto nitrocellulose membranes using a semidry transfer system (20 V, 45 min). Membranes were blocked in blocking buffer (5% w/v non-fat dried milk dissolved in TBST (50 mM Tris, 150 mM NaCl, pH 8.0, 0.1% v/v Tween-20)) at room temperature for 30 min. The primary antibody (rabbit anti-TnaA from AssayPro (cat. # 33517-05111) diluted 1:2000 in blocking buffer) was incubated overnight at 4°C. Following three 10-min washes with TBST, the secondary antibody (goat anti-rabbit, HRP-conjugated (Bio-Rad, cat. # 1705046), diluted 1:2000 in blocking buffer). Image acquisition was performed by adding 1 mL of mixed HRP substrate (Bio-Rad, cat. # 1705060) to the membrane for 1 min and then exposing the membrane to a cooled CCD camera (GE Healthcare ImageQuant LAS 500). Relative TnaA abundances were then estimated by densitometry using ImageJ.

### Proteomic analysis

Cultures of W3110 *E. coli* were grown at 30°C in five biological replicates for each strain, and harvested mid-log phase. Cells were lysed with lysozyme following resuspension. For the analysis of total (insoluble + soluble) protein content, lysed cells were adjusted to 8M urea and a brief ultrasonication (3x 10s) was performed to ensure complete protein solubilization. After a 2-hour digestion at 37°C with Lys-C, the solution was diluted to adjust urea to 2M; then trypsin was added and incubated overnight at 37°C. For the analysis of insoluble protein fractions, insoluble proteins were first isolated by high-speed centrifugation (20000g, 15 min, 4°C) following cell lysis. Pellets were then resuspended in 8M urea and solubilized by brief ultrasonication (3x 10s pulses), and digestion was performed similarly to the total cell lysates. To maximize digestion yields from the insoluble protein fractions, a second round of trypsin digestion was performed, followed by another ~16-hour incubation at 37°C.

In all cases, digests were desalted, resuspended in aqueous 2% acetonitrile + 0.05% trifluoroacetic acid. After loading onto a trapping microcolumn (Acclaim PepMap100 C18, 20 mm x 100 μm ID, 5 μm, Dionex), peptides were separated on a custom-packed nanocolumn (75 μm ID × 40 cm, 1.8 μm particles, Reprosil Pur, Dr. Maisch), with a flow rate of 250 nL/min and a gradient from 4% to 76% acetonitrile in water + 0.1% formic acid, over 140 min. Eluted peptides were analyzed on an Orbitrap Fusion Tribrid mass spectrometer (Thermo Fisher Scientific, Bremen, Germany) operated in data-dependent mode, controlled by Xcalibur software (version 3.0.63). Full survey scans were performed at a 120000 resolution, and a top speed precursor selection strategy was applied to maximize acquisition of peptide tandem MS spectra with a maximum cycle time of 3s. HCD fragmentation mode was used at a normalized collision energy of 32%, with a precursor isolation window of 1.6 m/z, and MS/MS spectra were acquired in the ion trap. Peptides selected for MS/MS were excluded from further fragmentation during 60s. Data collected by the mass spectrometer were processed for protein identification and quantification using MaxQuant version 1.5.3.30, using the Andromeda search engine set to search the UniProt database restricted to the *E. coli* (strain K12) proteome (UniProt proteome ID: UP000000625, number of sequences: 4306). Trypsin (cleavage after K,R) was used as the enzyme definition, allowing 2 missed cleavages. Carbamidomethylation of cysteine was specified as a fixed modification, while N-terminal acetylation of protein and oxidation of methionine were specified as variable modifications. The mass spectrometry proteomics data have been deposited to the ProteomeXchange Consortium via the PRIDE (Vizcaino, Csordas et al. 2016) partner repository with the dataset identifier PXD010014.

All data post-processing and statistical analyses were performed using custom Matlab scripts. iBAQ and LFQ data were used as the basis for quantification. Instead of only using raw iBAQ intensities, we took advantage of the additional normalization introduced by the LFQ method (Cox, Hein et al. 2014) to re-calculate “normalized iBAQs” by dividing LFQ intensities by the number of theoretically observable tryptic peptides, as specified in the original iBAQ publication (Schwanhausser, Busse et al. 2011). Since normalized iBAQs are proportional to protein molar quantities, protein mass fractions were obtained as 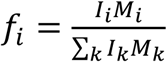 Where I_i_ is the normalized IBAQ intensity of protein i, M_i_ its molecular weight, and the index k runs over all identified proteins. Then, the corresponding micromolar quantities ci were derived using an estimated total intracellular protein concentration of C_T_ = 235 mg/mL (Zimmerman and Trach 1991, Ellis 2001): 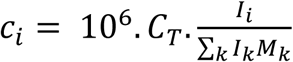

An additional normalization was then performed to correlate protein abundances in insoluble fractions to those in the total protein fractions. A list of 281 *bona* fide membrane proteins was obtained from the Uniprot database. These known, fully-insoluble proteins should therefore have the same mass fractions in the total and pellet mass fractions. For this equality to hold, log-log scatter plots of pellet *versus* total mass fractions (**Figure 2A**) were used to perform curve fitting procedures (separately for each *E. coli* strain) of the form Log P = αLog T + β, where P and T correspond to pellet and total mass fractions, respectively. The obtained normalization coefficients α and β were then applied to the pellet mass factions of all identified proteins.

Statistical analyses using our five biological replicates were then performed, first to determine which proteins were significantly quantified *(i.e*. had a mass fraction significantly larger tnan zero). This was done using *t* tests with *a post-hoc Benjamini-Hochberg* FDR-controlling procedure to account for multiple testing, at a FDR threshold of 0.01. Finally, significant differences in abundance or solubility between pairs of *E. coli* strains were determined using two-sample *t* tests followed by Benjamini-Hochberg procedures using an FDR cutoff of 0.05.

## References

Abravaya, K., M. P. Myers, S. P. Murphy and R. I. Morimoto (1992). “The human heat shock protein hsp70 interacts with HSF, the transcription factor that regulates heat shock gene expression.” Genes Dev 6(7): 1153–1164.

Angles, F., M. P. Castanie-Cornet, N. Slama, M. Dinclaux, A. M. Cirinesi, J. C. Portais, F. Letisse and P. Genevaux (2017). “Multilevel interaction of the DnaK/DnaJ(HSP70/HSP40) stress-responsive chaperone machine with the central metabolism.” Sci Rep 7: 41341.

Arike, L., K. Valgepea, L. Peil, R. Nahku, K. Adamberg and R. Vilu (2012). “Comparison and applications of label-free absolute proteome quantification methods on Escherichia coli.” J Proteomics 75(17): 5437–5448.

Ayrault, O., M. D. Godeny, C. Dillon, F. Zindy, P. Fitzgerald, M. F. Roussel and H. M. Beere (2009). “Inhibition of Hsp90 via 17-DMAG induces apoptosis in a p53-dependent manner to prevent medulloblastoma.” Proc Natl Acad Sci U S A 106(40): 17037–17042.

Azem, A., S. Diamant, M. Kessel, C. Weiss and P. Goloubinoff (1995). “The protein-folding activity of chaperonins correlates with the symmetric GroEL14(GroES7)2 heterooligomer.” Proc Natl Acad Sci U S A 92(26): 12021–12025.

Bachmann, B. J. (1972). “Pedigrees of some mutant strains of Escherichia coli K-12.” Bacteriol Rev 36(4): 525–557.

Balchin, D., M. Hayer-Hartl and F. U. Hartl (2016). “In vivo aspects of protein folding and quality control.” Science 353(6294): aac4354.

Bar-Lavan, Y., N. Shemesh, S. Dror, R. Ofir, E. Yeger-Lotem and A. Ben-Zvi (2016). “A Differentiation Transcription Factor Establishes Muscle-Specific Proteostasis in Caenorhabditis elegans.” PLoS Genet 12(12): e1006531.

Bardwell, J. C. and E. A. Craig (1988). “Ancient heat shock gene is dispensable.” J Bacteriol 170(7): 2977–2983.

Bukau, B. and G. C. Walker (1989). “Cellular defects caused by deletion of the Escherichia coli dnaK gene indicate roles for heat shock protein in normal metabolism.” J Bacteriol 171(5): 2337–2346.

Calloni, G., T. Chen, S. M. Schermann, H. C. Chang, P. Genevaux, F. Agostini, G. G. Tartaglia, M. Hayer-Hartl and F. U. Hartl (2012). “DnaK functions as a central hub in the E. coli chaperone network.” Cell Rep 1(3): 251–264.

Cox, J., M. Y. Hein, C. A. Luber, I. Paron, N. Nagaraj and M. Mann (2014). “Accurate proteome-wide label-free quantification by delayed normalization and maximal peptide ratio extraction, termed MaxLFQ.” Mol Cell Proteomics 13(9): 2513–2526.

Craig, R., J. P. Cortens and R. C. Beavis (2004). “Open source system for analyzing, validating, and storing protein identification data.” J Proteome Res 3(6): 1234–1242.

Dang, W., Y. H. Hu and L. Sun (2011). “HtpG is involved in the pathogenesis of Edwardsiella tarda.” Vet Microbiol 152(3-4): 394–400.

Deuerling, E. and B. Bukau (2004). “Chaperone-assisted folding of newly synthesized proteins in the cytosol.” Crit Rev Biochem Mol Biol 39(5-6): 261–277.

Diamant, S., A. P. Ben-Zvi, B. Bukau and P. Goloubinoff (2000). “Size-dependent disaggregation of stable protein aggregates by the DnaK chaperone machinery.” J Biol Chem 275(28): 21107–21113.

Echeverria, P. C. and D. Picard (2010). “Molecular chaperones, essential partners of steroid hormone receptors for activity and mobility.” Biochim Biophys Acta 1803(6): 641–649.

Ellis, R. J. (2001). “Macromolecular crowding: an important but neglected aspect of the intracellular environment.” Curr Opin Struct Biol 11(1): 114–119.

Fan, D., C. Liu, L. Liu, L. Zhu, F. Peng and Q. Zhou (2016). “Large-scale gene expression profiling reveals physiological response to deletion of chaperone dnaKJ in Escherichia coli.” Microbiol Res 186-187: 27–36.

Fierro-Monti, I., P. Echeverria, J. Racle, C. Hernandez, D. Picard and M. Quadroni (2013). “Dynamic impacts of the inhibition of the molecular chaperone Hsp90 on the T-cell proteome have implications for anti-cancer therapy.” PLoS One 8(11): e80425.

Fierro-Monti, I., J. Racle, C. Hernandez, P. Waridel, V. Hatzimanikatis and M. Quadroni (2013). “A novel pulse-chase SILAC strategy measures changes in protein decay and synthesis rates induced by perturbation of proteostasis with an Hsp90 inhibitor.” PLoS One 8(11): e80423.

Finka, A., A. F. Cuendet, F. J. Maathuis, Y. Saidi and P. Goloubinoff (2012). “Plasma membrane cyclic nucleotide gated calcium channels control land plant thermal sensing and acquired thermotolerance.” Plant Cell 24(8): 3333–3348.

Finka, A. and P. Goloubinoff (2013). “Proteomic data from human cell cultures refine mechanisms of chaperone-mediated protein homeostasis.” Cell Stress Chaperones 18(5): 591–605.

Finka, A., R. U. Mattoo and P. Goloubinoff (2011). “Meta-analysis of heat-and chemically upregulated chaperone genes in plant and human cells.” Cell Stress Chaperones 16(1): 15–31.

Finka, A., R. U. Mattoo and P. Goloubinoff (2016). “Experimental Milestones in the Discovery of Molecular Chaperones as Polypeptide Unfolding Enzymes.” Annu Rev Biochem 85: 715–742.

Finka, A., V. Sood, M. Quadroni, L. Rios Pde and P. Goloubinoff (2015). “Quantitative proteomics of heat-treated human cells show an across-the-board mild depletion of housekeeping proteins to massively accumulate few HSPs.” Cell Stress Chaperones 20(4): 605–620.

Finka, A., V. Sood, M. Quadroni, P. L. Rios and P. Goloubinoff (2015). “Quantitative proteomics of heat-treated human cells show an across-the-board mild depletion of housekeeping proteins to massively accumulate few HSPs.” Cell Stress Chaperones.

Flynn, J. M., P. Mishra and D. N. Bolon (2015). “Mechanistic Asymmetry in Hsp90 Dimers.” J Mol Biol 427(18): 2904–2911.

Franzosa, E. A., V. Albanese, J. Frydman, Y. Xia and A. J. McClellan (2011). “Heterozygous yeast deletion collection screens reveal essential targets of Hsp90.” PLoS One 6(11): e28211.

Garcie, C., S. Tronnet, A. Garenaux, A. J. McCarthy, A. O. Brachmann, M. Penary, S. Houle, J. P. Nougayrede, J. Piel, P. W. Taylor, C. M. Dozois, P. Genevaux, E. Oswald and P. Martin (2016). “The Bacterial Stress-Responsive Hsp90 Chaperone (HtpG) Is Required for the Production of the Genotoxin Colibactin and the Siderophore Yersiniabactin in Escherichia coli.” J Infect Dis 214(6): 916–924.

Gat-Yablonski, G., A. Finka, G. Pinto, M. Quadroni, B. Shtaif and P. Goloubinoff (2016). “Quantitative proteomics of rat livers shows that unrestricted feeding is stressful for proteostasis with implications on life span.” Aging (Albany NY) 8(8): 1735–1758.

Genest, O., J. R. Hoskins, J. L. Camberg, S. M. Doyle and S. Wickner (2011). “Heat shock protein 90 from Escherichia coli collaborates with the DnaK chaperone system in client protein remodeling.” Proc Natl Acad Sci U S A 108(20): 8206–8211.

Genest, O., J. R. Hoskins, A. N. Kravats, S. M. Doyle and S. Wickner (2015). “Hsp70 and Hsp90 of E. coli Directly Interact for Collaboration in Protein Remodeling.” J Mol Biol 427(24): 3877–3889.

Genest, O., M. Reidy, T. O. Street, J. R. Hoskins, J. L. Camberg, D. A. Agard, D. C. Masison and S. Wickner (2013). “Uncovering a region of heat shock protein 90 important for client binding in E. coli and chaperone function in yeast.” Mol Cell 49(3): 464–473.

Genevaux, P., F. Keppel, F. Schwager, P. S. Langendijk-Genevaux, F. U. Hartl and C. Georgopoulos (2004). “In vivo analysis of the overlapping functions of DnaK and trigger factor.” EMBO Rep 5(2): 195–200.

Goloubinoff, P., A. Mogk, A. P. Zvi, T. Tomoyasu and B. Bukau (1999). “Sequential mechanism of solubilization and refolding of stable protein aggregates by a bichaperone network.” Proc Natl Acad Sci U S A 96(24): 13732–13737.

Goloubinoff, P., A. S. Sassi, B. Fauvet, A. Barducci and P. De Los Rios (2018). “Chaperones convert the energy from ATP into the nonequilibrium stabilization of native proteins.” Nature Chemical Biology.

Grudniak, A. M., K. Pawlak, K. Bartosik and K. I. Wolska (2013). “Physiological consequences of mutations in the htpG heat shock gene of Escherichia coli.” Mutat Res 745-746: 1–5.

Hartl, F. U. (2017). “Protein Misfolding Diseases.” Annu Rev Biochem 86: 21–26.

Hinault, M. P., A. Farina-Henriquez-Cuendet and P. Goloubinoff (2011). “Molecular chaperones and associated cellular clearance mechanisms against toxic protein conformers in Parkinson’s disease.” Neurodegener Dis 8(6): 397–412.

Honore, F. A., V. Mejean and O. Genest (2017). “Hsp90 Is Essential under Heat Stress in the Bacterium Shewanella oneidensis.” Cell Rep 19(4): 680–687.

Hossain, M. M. and H. Nakamoto (2003). “Role for the cyanobacterial HtpG in protection from oxidative stress.” Curr Microbiol 46(1): 70–76.

Jaattela, M., D. Wissing, K. Kokholm, T. Kallunki and M. Egeblad (1998). “Hsp70 exerts its anti-apoptotic function downstream of caspase-3-like proteases.” EMBO J 17(21): 6124–6134.

Katayama, Y., S. Gottesman, J. Pumphrey, S. Rudikoff, W. P. Clark and M. R. Maurizi (1988). “The two-component, ATP-dependent Clp protease of Escherichia coli. Purification, cloning, and mutational analysis of the ATP-binding component.” J Biol Chem 263(29): 15226–15236.

Kitagawa, M., T. Ara, M. Arifuzzaman, T. Ioka-Nakamichi, E. Inamoto, H. Toyonaga and H. Mori (2005). “Complete set of ORF clones of Escherichia coli ASKA library (a complete set of E. coli K-12 ORF archive): unique resources for biological research.” DNA Res 12(5): 291–299.

Kitagawa, M., M. Miyakawa, Y. Matsumura and T. Tsuchido (2002). “Escherichia coli small heat shock proteins, IbpA and IbpB, protect enzymes from inactivation by heat and oxidants.” Eur J Biochem 269(12): 2907–2917.

Kravats, A. N., S. M. Doyle, J. R. Hoskins, O. Genest, E. Doody and S. Wickner (2017). “Interaction of E. coli Hsp90 with DnaK Involves the DnaJ Binding Region of DnaK.” J Mol Biol 429(6): 858–872.

Krug, K., A. Carpy, G. Behrends, K. Matic, N. C. Soares and B. Macek (2013). “Deep coverage of the Escherichia coli proteome enables the assessment of false discovery rates in simple proteogenomic experiments.” Mol Cell Proteomics 12(11): 3420–3430.

Krukenberg, K. A., T. O. Street, L. A. Lavery and D. A. Agard (2011). “Conformational dynamics of the molecular chaperone Hsp90.” Q Rev Biophys 44(2): 229–255.

Lackie, R. E., A. Maciejewski, V. G. Ostapchenko, J. Marques-Lopes, W. Y. Choy, M. L. Duennwald, V. F. Prado and M. A. M. Prado (2017). “The Hsp70/Hsp90 Chaperone Machinery in Neurodegenerative Diseases.” Front Neurosci 11: 254.

Lanneau, D., A. de Thonel, S. Maurel, C. Didelot and C. Garrido (2007). “Apoptosis versus cell differentiation: role of heat shock proteins HSP90, HSP70 and HSP27.” Prion 1(1): 53–60.

Lashuel, H. A. and P. T. Lansbury, Jr. (2006). “Are amyloid diseases caused by protein aggregates that mimic bacterial pore-forming toxins?” Q Rev Biophys 39(2): 167–201.

Lee, J. H., J. Gao, P. A. Kosinski, S. J. Elliman, T. E. Hughes, J. Gromada and D. M. Kemp (2013). “Heat shock protein 90 (HSP90) inhibitors activate the heat shock factor 1 (HSF1) stress response pathway and improve glucose regulation in diabetic mice.” Biochem Biophys Res Commun 430(3): 1109–1113.

Li, J., J. Soroka and J. Buchner (2012). “The Hsp90 chaperone machinery: conformational dynamics and regulation by co-chaperones.” Biochim Biophys Acta 1823(3): 624–635.

Lim, B., R. Miyazaki, S. Neher, D. A. Siegele, K. Ito, P. Walter, Y. Akiyama, T. Yura and C. A. Gross (2013). “Heat shock transcription factor sigma32 co-opts the signal recognition particle to regulate protein homeostasis in E. coli.” PLoS Biol 11(12): e1001735.

Mayer, M. P. and L. Le Breton (2015). “Hsp90: breaking the symmetry.” Mol Cell 58(1): 8–20.

Mogk, A., T. Tomoyasu, P. Goloubinoff, S. Rudiger, D. Roder, H. Langen and B. Bukau (1999). “Identification of thermolabile Escherichia coli proteins: prevention and reversion of aggregation by DnaK and ClpB.” EMBO J 18(24): 6934–6949.

Motojima-Miyazaki, Y., M. Yoshida and F. Motojima (2010). “Ribosomal protein L2 associates with E. coli HtpG and activates its ATPase activity.” Biochem Biophys Res Commun 400(2): 241–245.

Multhoff, G. (1997). “Heat shock protein 72 (HSP72), a hyperthermia-inducible immunogenic determinant on leukemic K562 and Ewing’s sarcoma cells.” Int J Hyperthermia 13(1): 39–48.

Nakamoto, H., K. Fujita, A. Ohtaki, S. Watanabe, S. Narumi, T. Maruyama, E. Suenaga, T. S. Misono, P. K. Kumar, P. Goloubinoff and H. Yoshikawa (2014). “Physical interaction between bacterial heat shock protein (Hsp) 90 and Hsp70 chaperones mediates their cooperative action to refold denatured proteins.” J Biol Chem 289(9): 6110–6119.

Neckers, L., A. Kern and S. Tsutsumi (2007). “Hsp90 inhibitors disrupt mitochondrial homeostasis in cancer cells.” Chem Biol 14(11): 1204–1206.

Pratt, W. B., Y. Morishima, H. M. Peng and Y. Osawa (2010). “Proposal for a role of the Hsp90/Hsp70-based chaperone machinery in making triage decisions when proteins undergo oxidative and toxic damage.” Exp Biol Med (Maywood) 235(3): 278–289.

Press, M. O., H. Li, N. Creanza, G. Kramer, C. Queitsch, V. Sourjik and E. Borenstein (2013). “Genome-scale co-evolutionary inference identifies functions and clients of bacterial Hsp90.” PLoS Genet 9(7): e1003631.

Sato, T., S. Minagawa, E. Kojima, N. Okamoto and H. Nakamoto (2010). “HtpG, the prokaryotic homologue of Hsp90, stabilizes a phycobilisome protein in the cyanobacterium Synechococcus elongatus PCC 7942.” Mol Microbiol 76(3): 576–589.

Schmidt, A., K. Kochanowski, S. Vedelaar, E. Ahrne, B. Volkmer, L. Callipo, K. Knoops, M. Bauer, R. Aebersold and M. Heinemann (2016). “The quantitative and condition-dependent Escherichia coli proteome.” Nat Biotechnol 34(1): 104–110.

Schramm, F. D., K. Heinrich, M. Thuring, J. Bernhardt and K. Jonas (2017). “An essential regulatory function of the DnaK chaperone dictates the decision between proliferation and maintenance in Caulobacter crescentus.” PLoS Genet 13(12): e1007148.

Schwanhausser, B., D. Busse, N. Li, G. Dittmar, J. Schuchhardt, J. Wolf, W. Chen and M. Selbach (2011). “Global quantification of mammalian gene expression control.” Nature 473(7347): 337–342.

Sharma, S. K., P. De Los Rios and P. Goloubinoff (2011). “Probing the different chaperone activities of the bacterial HSP70-HSP40 system using a thermolabile luciferase substrate.” Proteins 79(6): 1991–1998.

Sherman, M. Y. and A. L. Goldberg (1996). “Involvement of molecular chaperones in intracellular protein breakdown.” EXS 77: 57–78.

Shi, W., Y. Zhou, J. Wild, J. Adler and C. A. Gross (1992). “DnaK, DnaJ, and GrpE are required for flagellum synthesis in Escherichia coli.” J Bacteriol 174(19): 6256–6263.

Shiau, A. K., S. F. Harris, D. R. Southworth and D. A. Agard (2006). “Structural Analysis of E. coli hsp90 reveals dramatic nucleotide-dependent conformational rearrangements.” Cell 127(2): 329–340.

Stefani, M. and C. M. Dobson (2003). “Protein aggregation and aggregate toxicity: new insights into protein folding, misfolding diseases and biological evolution.” J Mol Med (Berl) 81(11): 678–699.

Street, T. O., X. Zeng, R. Pellarin, M. Bonomi, A. Sali, M. J. Kelly, F. Chu and D. A. Agard (2014). “Elucidating the mechanism of substrate recognition by the bacterial Hsp90 molecular chaperone.” J Mol Biol 426(12): 2393–2404.

Taipale, M., D. F. Jarosz and S. Lindquist (2010). “HSP90 at the hub of protein homeostasis: emerging mechanistic insights.” Nat Rev Mol Cell Biol 11(7): 515–528.

Tanaka, N. and H. Nakamoto (1999). “HtpG is essential for the thermal stress management in cyanobacteria.” FEBS Lett 458(2): 117–123.

Thomas, J. G. and F. Baneyx (1998). “Roles of the Escherichia coli small heat shock proteins IbpA and IbpB in thermal stress management: comparison with ClpA, ClpB, and HtpG In vivo.” J Bacteriol 180(19): 5165–5172.

Thomas, J. G. and F. Baneyx (2000). “ClpB and HtpG facilitate de novo protein folding in stressed Escherichia coli cells.” Mol Microbiol 36(6): 1360–1370.

Valgepea, K., K. Adamberg, R. Nahku, P. J. Lahtvee, L. Arike and R. Vilu (2010). “Systems biology approach reveals that overflow metabolism of acetate in Escherichia coli is triggered by carbon catabolite repression of acetyl-CoA synthetase.” BMC Syst Biol 4: 166.

Vartholomaiou, E., P. C. Echeverria and D. Picard (2016). “Unusual Suspects in the Twilight Zone Between the Hsp90 Interactome and Carcinogenesis.” Adv Cancer Res 129: 1–30.

Verbrugghe, E., A. Van Parys, B. Leyman, F. Boyen, F. Haesebrouck and F. Pasmans (2015). “HtpG contributes to Salmonella Typhimurium intestinal persistence in pigs.” Vet Res 46: 118.

Vivien, E., S. Megessier, I. Pieretti, S. Cociancich, R. Frutos, D. W. Gabriel, P. C. Rott and M. Royer (2005). “Xanthomonas albilineans HtpG is required for biosynthesis of the antibiotic and phytotoxin albicidin.” FEMS Microbiol Lett 251(1): 81–89.

Vizcaino, J. A., A. Csordas, N. del-Toro, J. A. Dianes, J. Griss, I. Lavidas, G. Mayer, Y. Perez-Riverol, F. Reisinger, T. Ternent, Q. W. Xu, R. Wang and H. Hermjakob (2016). “2016 update of the PRIDE database and its related tools.” Nucleic Acids Res 44(D1): D447–456.

Weiss, Y. G., Z. Bromberg, N. Raj, J. Raphael, P. Goloubinoff, Y. Ben-Neriah and C. S. Deutschman (2007). “Enhanced heat shock protein 70 expression alters proteasomal degradation of IkappaB kinase in experimental acute respiratory distress syndrome.” Crit Care Med 35(9): 2128–2138.

Wiech, H., J. Buchner, R. Zimmermann and U. Jakob (1992). “Hsp90 chaperones protein folding in vitro.” Nature 358(6382): 169–170.

Yosef, I., M. G. Goren, R. Kiro, R. Edgar and U. Qimron (2011). “High-temperature protein G is essential for activity of the Escherichia coli clustered regularly interspaced short palindromic repeats (CRISPR)/Cas system.” Proceedings of the National Academy of Sciences of the United States of America 108(50): 20136–20141.

Yosef, I., M. G. Goren, R. Kiro, R. Edgar and U. Qimron (2011). “High-temperature protein G is essential for activity of the Escherichia coli clustered regularly interspaced short palindromic repeats (CRISPR)/Cas system.” Proc Natl Acad Sci U S A 108(50): 20136–20141.

Zimmerman, S. B. and S. O. Trach (1991). “Estimation of macromolecule concentrations and excluded volume effects for the cytoplasm of Escherichia coli.” J Mol Biol 222(3): 599–620.

